# Temperature and *skn-1* determine whether developmental *daf-2* knockdown extends *C. elegans* lifespan

**DOI:** 10.64898/2026.11.18.767509

**Authors:** Sara D. Irish, Annabel Kimberley, Simone Immler, Alexei A. Maklakov

## Abstract

The evolutionarily conserved insulin/IGF-1 signalling pathway (IIS) shapes life-history traits largely through downstream transcription factors including SKN-1. Reduced IIS (rIIS) in adulthood robustly extends *Caenorhabditis elegans* lifespan, yet developmental rIIS has long been considered unable to extend lifespan and costly to fitness, a conclusion drawn almost entirely from experiments at 25°C, near the species’ upper thermal limit. Here we knocked down *daf-2* during development in wild-type and *skn-1* loss-of-function nematodes across constant and fluctuating temperatures. Contrary to the prevailing view, developmental *daf-2* knockdown extended lifespan at constant 15°C in both backgrounds, but not at 25°C, revealing that its late-life benefit is strongly temperature-dependent and, under constant conditions, independent of *skn-1*. Under daily 15–25°C cycling, developmental *daf-2* knockdown again extended lifespan, but whether this required *skn-1* was itself determined by the temperature experienced at the start of development. Lifespan extension was accompanied by condition-specific reproductive costs. Together, our results identify a developmental window in which IIS, *skn-1* and temperature interact to set adult ageing trajectories and show that the thermal environment determines whether reducing IIS during development extends lifespan.

## 1. Introduction

Manipulation of the highly evolutionarily conserved insulin-signalling sensing pathway (IIS) is a well-established intervention to influence life history traits across a range of model organisms (Clancy et al., 2001; Fabrizio et al., 2001; Holzenberger et al., 2003). In *Caenorhabditis elegans* nematodes, the *daf-2* receptor controls IIS levels, and mutations or manipulations that decrease *daf-2* activity can dramatically extend lifespan (Dillin et al., 2002; Kenyon, 2001; Kenyon et al., 1993). Reduced IIS (rIIS) has also been implicated in improved somatic maintenance (Murphy et al., 2003; Taylor and Dillin, 2011), stress resistance (Carlsson et al., 2021) and anti-bacterial (Garsin et al., 2003) and anti-viral immunity (Duxbury et al., 2024), traits that are critical to survival.

Despite extensive work on IIS in adulthood, the role of IIS regulation during development in shaping environmental responses and long-term life-history trajectories remains comparatively poorly understood. Early work demonstrated that *daf-2* knockdown during development does not extend lifespan and is detrimental to fitness through reproductive delays (Dillin et al., 2002), leading to the prevailing assumption that reduced IIS is maladaptive in early life. In adulthood, rIIS extends lifespan without fitness costs and can even improve Darwinian fitness of offspring (Lind et al., 2019). Nevertheless, little attention has been given to the effects of developmental IIS regulation on life-history traits outside of mutants. Moreover, the experiment specifically testing developmental rIIS used *daf-2* RNAi knockdown initiated at hatching and conducted at 25°C (Dillin et al. 2002), a temperature approaching the upper thermal limit of *C. elegans,* and one that populations are unlikely to experience frequently in nature (Félix and Duveau, 2012). This calls into question whether the detrimental fitness effects of developmental rIIS may be a laboratory artefact, especially considering the importance of the role of IIS responses to environmental cues (Regan et al., 2020).

Importantly, IIS is likely tightly regulated throughout stages of larval development and plays a central role in mediating developmental plasticity in response to environmental cues (Kaplan et al., 2019). In particular, IIS controls entry into dauer diapause, a non-feeding, stress-resistant developmental stage that allows *C. elegans* to survive adverse conditions such as crowding, food scarcity, and thermal stress (Gems et al., 1998; Golden and Riddle, 1984; Kimura et al., 1997). In nature, *C. elegans* populations experience boom-and-bust dynamics driven by ephemeral food sources, and dauer entry not only promotes persistence in these conditions, but also facilitates dispersal to new food sources (Schulenburg and Félix, 2017). Although rIIS during development may confer benefits under some conditions, selection acting on early-life IIS regulation is likely constrained by the requirement to maintain developmental responsiveness to environmental stressors.

Environmental temperature during development is itself a major determinant in shaping *C. elegans* life history. Relatively small changes in temperature (± 5°C) have been shown to substantially alter gene expression (Zhang et al., 2015), developmental timing (Klass, 1977), and lifespan (Xiao et al., 2013). While temperature effects on lifespan have often been attributed to changes in metabolic rate, accumulating evidence suggests that temperature also modulates ageing through transcriptional regulation downstream of IIS. In particular, the IIS-associated transcription factors DAF-16, HSF-1, and SKN-1 have been implicated in mediating temperature-dependent effects on stress resistance and longevity (Chiang et al., 2012; Ewald et al., 2015; Galbadage and Hartman, 2008; Xiao et al., 2013).

SKN-1 is known to exert effects on development, stress resistance, and protein homeostasis (Blackwell et al., 2015; Tullet et al., 2008; Turner and Curran, 2025), with critical roles during embryogenesis (An and Blackwell, 2003; Bowerman et al., 1992), and post-embryonic somatic maintenance (Blackwell et al., 2015; Li et al., 2011). Notably, both insufficient and excessive activity of SKN-1 can negatively impact lifespan (Lynn et al., 2015; Zhang et al., 2015). This indicates that SKN-1 plays important roles across life-history traits from embryogenesis to ageing, yet how IIS and SKN-1 interact with early-life environmental temperatures remains largely unknown.

Improving our understanding of how IIS interacts with environmental challenges, such as temperature stress, is critical for elucidating the role of this evolutionarily conserved pathway in life-history regulation. Here, we tested how rIIS*, skn-1,* age, and temperature interact in a series of experiments using *daf-2* RNAi knockdown during larval development in *C. elegans skn-1* loss-of-function (mg570) mutants, which exhibit limited somatic stress resistance (Lehrbach and Ruvkun, 2019). We found that there is a critical period during development in which IIS, *skn-1*, and temperature interact to shape ageing trajectories, indicating specific early-life gene expression requirements for regulating life-history. Although *daf-2* RNAi extended lifespan across larval temperature cycles, environmental variability exposed fitness costs through delayed reproduction, demonstrating a potential developmental constraint that limits optimal IIS regulation for improved late-life survival.

## 2. Methods

### 2.1. Experimental model

Strains of *Caenorhabditis elegans* nematodes were obtained from the *Caenorhabditis* Genetics Centre, funded by NIH Office of Research Infrastructure Programs (P40 OD010440). Nematodes stored at -70°C were thawed and once populations began producing eggs, they were bleached using a solution of NaOCl and NaOH to remove contaminants. Populations were maintained at 20°C for at least two generations before use on 90 mm Nematode Growth Medium (NGM) plates containing 10 μg ml^-1^ of Nystatin and 100 μg ml^-1^ of ampicillin. Their maintenance diet consisted of *Escherichia coli* OP50-1 (pUC4 K). Age-synchronised populations were produced using short-duration egg lays, in which peak reproductive adults were allowed to lay eggs for a maximum of two hours. These eggs are then used to produce experimental populations.

To induce *daf-2* RNAi knockdown, nematodes were fed *E. coli* HT115 (DE3) containing a pL4440 vector which contained an insert expressing doubt-stranded RNA corresponding to the *daf-2* gene. Controls were fed an *E. coli* clone containing the pL4440 vector without the insert (“empty vector”, or ev, hereon). These were seeded onto NGM plates containing 10 μg ml^-1^ of nystatin and 50 μg ml^-1^ ampicillin and 1 mM IPTG, required for *daf-2* RNAi insert transcription.

### 2.2. Lifespan and reproduction assays

Nematodes for all assays were fed *daf-2* RNAi bacteria (or the ev control bacteria) upon hatching and were moved onto OP50-1 *E. coli* upon reaching sexual maturity (late L4 larval stage). For lifespan assays, nematodes were housed in groups of ten on 35-mm NGM plates to prevent starvation and overpopulation. During the reproductive period, nematodes were transferred to fresh plates daily to further prevent starvation. When reproduction ended, transfers occurred every other day to minimise stress. Daily mortality was determined by lack of response to touch. Deaths due to human error or unnatural causes were recorded and excluded from analysis. Additionally, matricides, where eggs hatch inside the mother leading to death, were recorded to analyse both with and without them censored. To test the effect of developmental *daf-2* knockdown on reproduction, nematodes were kept individually on plates upon reaching sexual maturity and transferred every 24-hours onto new plates until reproduction ended. Offspring from each day were killed and counted upon reaching approximately the L4 larval stage.

First, we tested the effects of *daf-2* RNAi during development in *skn-1* mutants on lifespan and reproduction at our standard laboratory temperature 20°C. To examine the interaction between temperature, IIS, and developmental timing, we used thermal regimes spanning 15°C and 25°C, which represent the lower and upper bounds of temperatures commonly used for laboratory maintenance of *C. elegans* (Stiernagle, 2006). For thermocycling assays, incubators set to constant 15°C or 25°C were used, and nematodes were transferred between temperatures at the same time each day. Two lifelong thermocycling regimes were employed, differing in the initial temperature experienced from the egg stage (15°C or 25°C), after which nematodes alternated daily between temperatures (see Fig. 1A).

**Figure 1.**
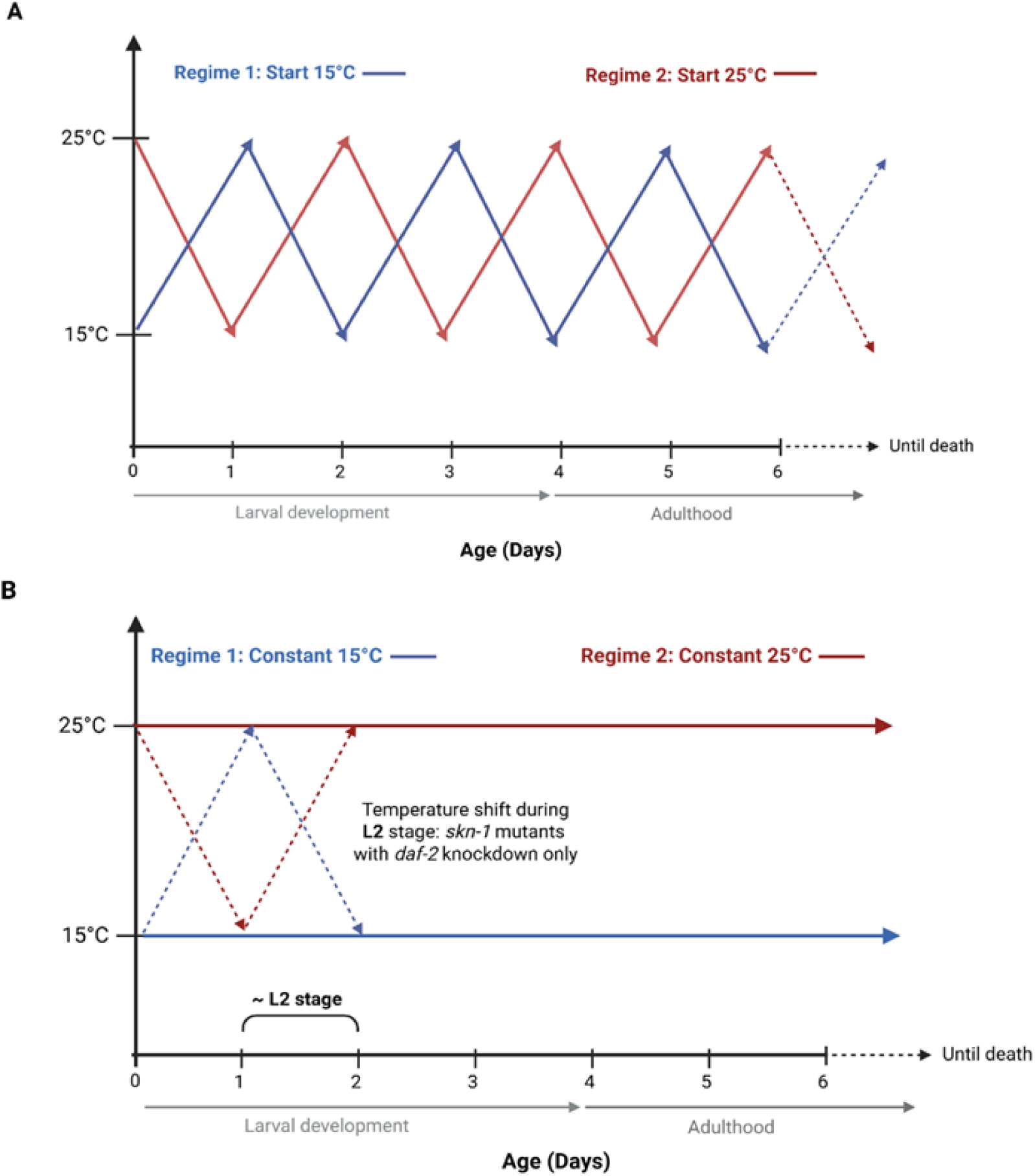
Schematic of thermocycling regimes employed from the egg stage in N2 and *skn-1* mutants with and without *daf-2* RNAi knockdown in development. In **(A)**, daily thermocycling between 15 and 25°C took place over the whole lifespan. In **(B)**, constant temperature treatment of 15 or 25°C was maintained lifelong, except for in *skn-1* mutants with *daf-2* knockdown, which were shifted for 24 hours to the opposite temperature during the L2 larval stage. All experimental individuals were selected as eggs, and immediately put into their respective starting temperature treatments. Diagonal lines do not represent gradual temperature shifts, they denote only the direction of the change on a given day. This figure was developed with BioRender.com.

Previous work has identified the L1 and L2 larval stages as critical periods for temperature-dependent regulation of stress response pathways in *C. elegans* (An and Blackwell, 2003; Zhang et al., 2015). To specifically target this developmental window, we conducted an additional assay in which N2 and *skn-1* mutants, with or without *daf-2* RNAi, were maintained at constant 15°C or 25°C, with *skn-1* mutants with developmental *daf-2* knockdown subjected to a single 24-hour thermocycle restricted to the L2 stage (see Fig. 1B). To minimise the effects of the ambient laboratory temperature (20 to 23°C) on the experiments, plates were kept out for a maximum of 10 minutes during transfers. All assays were blinded with respect to treatment.

### 2.3. Statistical analysis

All plotting and analysis were performed in R version 4.4.1 (R Core Team, 2022). Survival analyses were performed with Cox mixed effects models from the *coxme* package (Therneau, 2024) to compare hazard ratios between strains and treatments. For the thermocycling assay data, strain and treatment were fit as fixed effects, along with their interactions, and Plate ID was fit as a random intercept. Where interactions were non-significant, they were removed from the model. For the constant temperature experiment, with an additional developmental thermocycle for *skn-1* mutants with *daf-2* knockdown, survival analyses were performed within strain due to the unbalanced design, with Treatment as a fixed factor and Plate ID as a random intercept. Survival analyses were performed on all data including and excluding matricides, where death occurs due to eggs hatching inside the nematode. A phenomenon where ruptures occur and the intestines are expelled from the body, referred to as ‘Explosions’ in our data, were also excluded in all lifespan analyses (see Table S1 for summary of all lifespan experiments, including sample sizes and censoring). Additionally, any nematode losses due to human error or ‘walling,’ where individuals climb the walls of the agar plate and die due to desiccation, were also censored.

Age-specific reproduction data were analysed using zero-inflated negative binomial or Poisson generalised linear mixed effects models with the *glmmTMB* package (Brooks et al., 2024). Lifetime reproductive success was calculated as the total number of offspring produced per individual over the assay time and was analysed using general linear models. Alternatively, a Gamma family generalised linear model was used to analyse when the data were negatively skewed. To calculate individual fitness (λ), the *popbio* package v. 2.7 (Stubben and Milligan, 2007) was used to construct individual-based age-structured Leslie matrices (Leslie, 1945) which were used to calculate rate-sensitive fitness. Two days, the approximate time to reach sexual maturation for all treatments and temperature combinations, were added to the fertility schedule to represent development time (from egg until maturity). These data were also analysed using general linear models. Model diagnostics were performed with the *DHARMa* package (Hartig and Lohse, 2022), with model fits being chosen based on the most acceptable dispersion and zero-inflation parameters and comparison with Akaike information criterion.

## 3. Results

### 3.1. IIS downregulation does not extend lifespan in skn-1 mutants in 20°C

At 20°C, developmental *daf-2* knockdown did not extend lifespan in *skn-1* mutants regardless of whether matricides were excluded or included as natural deaths (Mixed effects Cox proportional hazards (COXME) excluding matricides: Χ_1_^2^ = 0.523, *p* = 0.466, Fig. 2A, matricides included: Χ_1_^2^ = 0.527, *p* = 0.468). Knocking down *daf-2* in *skn-1* mutants delayed age-specific reproduction (zero-inflated Poisson generalised linear mixed model (ZIP-GLMM): Treatment*Day^2^: Χ_1_^2^ = 7.101, *p* = 0.008, Table S2, Fig. 2B), confirmed by a 7 percent reduction in rate-sensitive fitness (LM: F_1,57_ = 4.69, *p* = 0.035, Fig. 2D), and lifetime reproductive success was reduced by 15 percent (LRS; Gamma generalised linear model (G-GLM): LR *Χ_1_^2^* = 4.100, *p* = 0.048, Fig. 2C).

**Figure 2.**
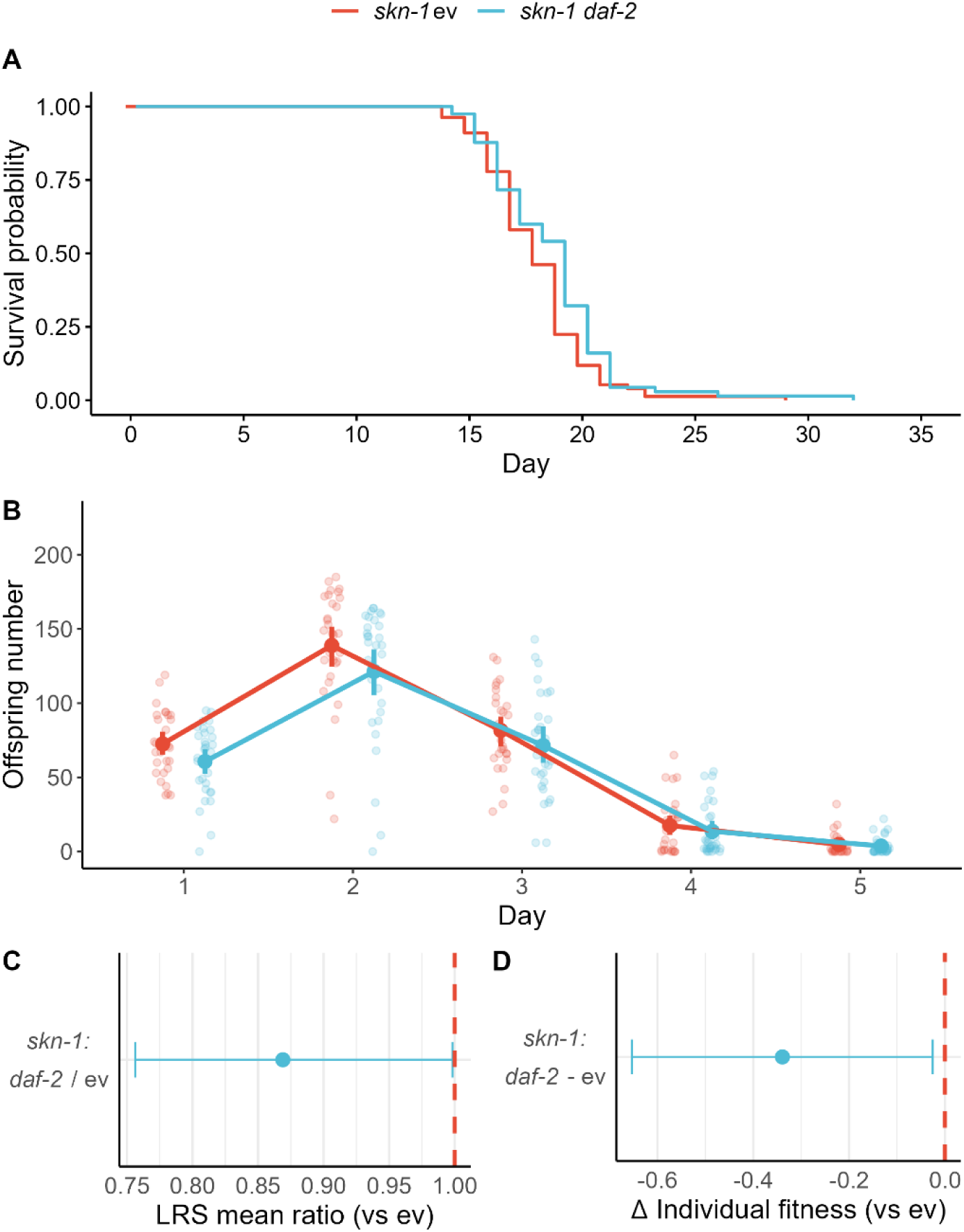
Developmental *daf-2* knockdown does not extend lifespan or affect age-specific reproduction in *skn-1* mutants at 20°C. (**A**) survival probability (n = 94 per treatment) shown with Kaplan-Meier survival curves, and (**B**) age-specific reproduction (n_ev_ = 30, n_daf-2_ = 31) with coloured points representing means with 95% confidence intervals (CIs)) and transparent points showing raw data. In (**C**) differences in lifetime reproductive success (LRS) shown by the mean ratio (with 95% CIs) calculated from a Gamma generalised linear model, with the red dotted line representing control baseline set to 1. In (**D**) the mean difference (with 95% CIs) in individual fitness calculated from a linear model is shown, with the red dotted line representing control baseline set to 0.

### 3.2. Reduced IIS extends lifespan under daily thermocycling in a partially skn-1 dependent manner

We found that *daf-2* knockdown during the developmental period extended lifespan only in the non-mutant N2 strain under thermocycling starting at 15°C (GLMM: matricides censored: Strain*Treatment: *Χ_1_^2^* = 61.562, *p* < 0.0001; Estimated marginal means (EMM) contrast: N2 *daf-2* – ev: -2.10, *p* < 0.001, Fig. 3A & B, Table S3). When the thermocycle began at 25°C, however, lifespan was extended by developmental *daf-2* knockdown in both strains (GLMM with matricides censored: Strain*Treatment: *Χ_1_^2^* = 7.324, *p* = 0.007; EMM contrast: N2 *daf-2* – ev: -2.07, p < 0.001; *skn-1 daf-2* – ev: -2.73, *p* < 0.0001, see Fig. 3C & D, Table S3). Including matricides in the analyses had no effect on outcomes (see Table S3).

**Figure 3.**
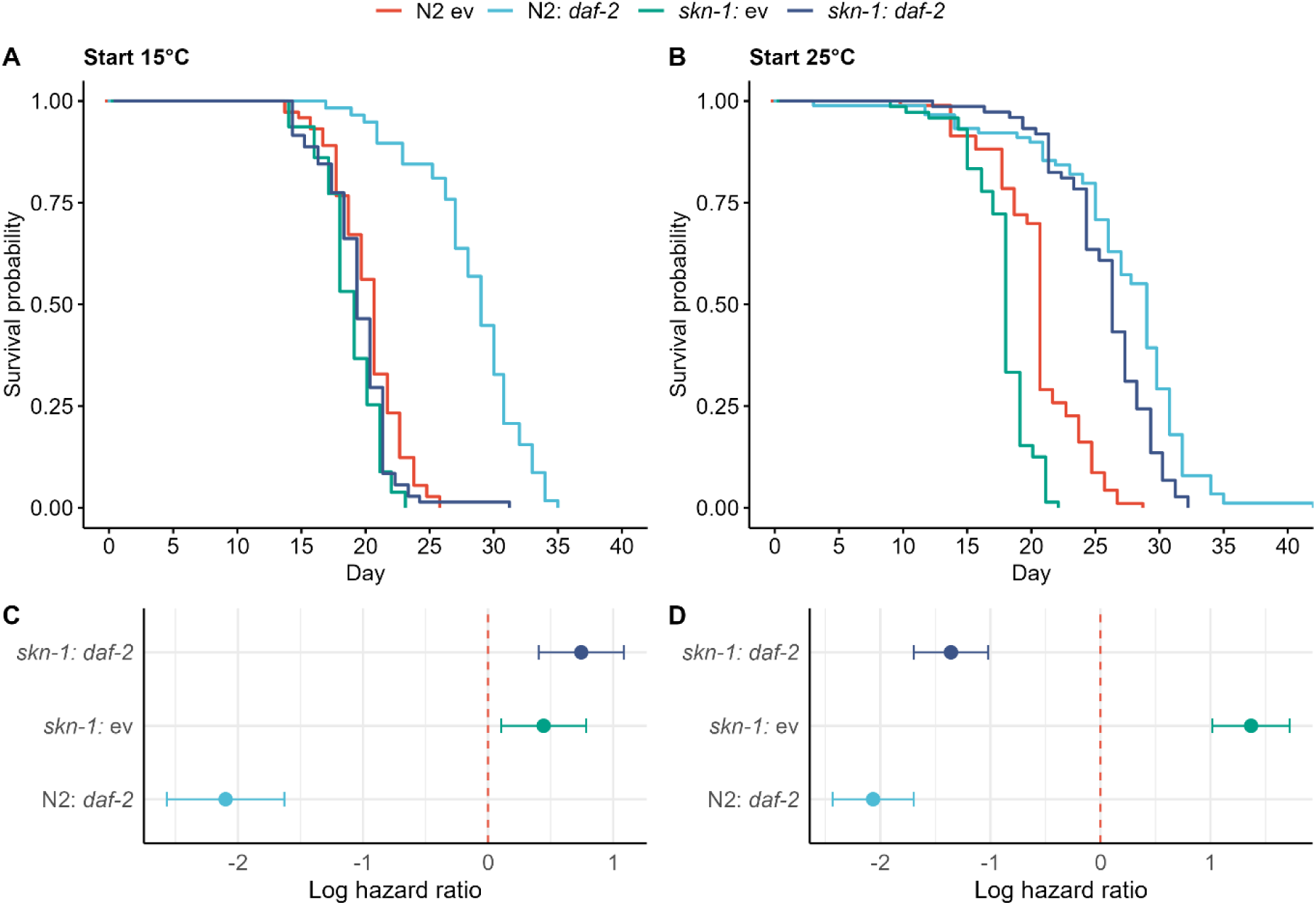
Developmental *daf-2* knockdown mediated lifespan extension is dependent on *skn-1* function when starting development at 15°C, but not at 25°C under a 24-hour thermocycle. **(A, B)** Survival probability shown with Kaplan-Meier survival curves and **(C, D)** corresponding log hazard ratios under 24-hr rapid thermocycle starting at 15°C (**A, C,** n > 57 per treatment x strain) and 25°C (**B, D**; n > 71 per treatment x strain) following *daf-2* knockdown in N2 Bristol strain nematodes and *skn-1* loss of function mutants. Dotted lines represent baseline log hazard ratio set to zero for controls (N2 or *skn-1* with ev treatment). Log hazard ratios represented by means and 95% confidence intervals.

### 3.3. Downregulated IIS under thermocycling reduces fitness

Starting thermocycling regimes at 15°C carried a cost to early reproduction following developmental *daf-2* knockdown in N2s, but not *skn-1* mutants (GLMM: Strain*Treatment*Day: *Χ_1_^2^* = 13.504, *p* = 0.0002, see Fig. 4A, Table S4). Confirming this, *daf-2* RNAi treatment reduced rate-sensitive individual fitness in N2s by 10 percent (LM: F_1,93_ = 5.414, *p* = 0.022, Fig. 4C), but this did not translate to a significant effect on total reproductive output in either N2 or *skn-1* nematodes (LM: F_1,115_ = 0.378, *p* = 0.540, Fig. 4B). However, reproductive costs to developmental *daf-2* knockdown in *skn-1* mutants appear to occur under thermocycling starting at 25°C through delays in early reproduction (GLMM: Treatment*Day: *Χ_1_^2^* = 49.505, *p* = <0.0001; Strain*Day: *Χ_1_^2^* = 11.249, *p* = 0.001, see Fig. 4D, Table S4), which caused a 12 percent decrease in individual fitness (LM: Strain*Treatment: F_1,98_ = 7.093, *p* = 0.009, Fig. 4F). There was also an overall cost to LRS in *skn-1* mutants regardless of treatment (LM: F_1,117_ = 6.069, *p* = 0.015, Fig. 4E, Table S5).

**Figure 4.**
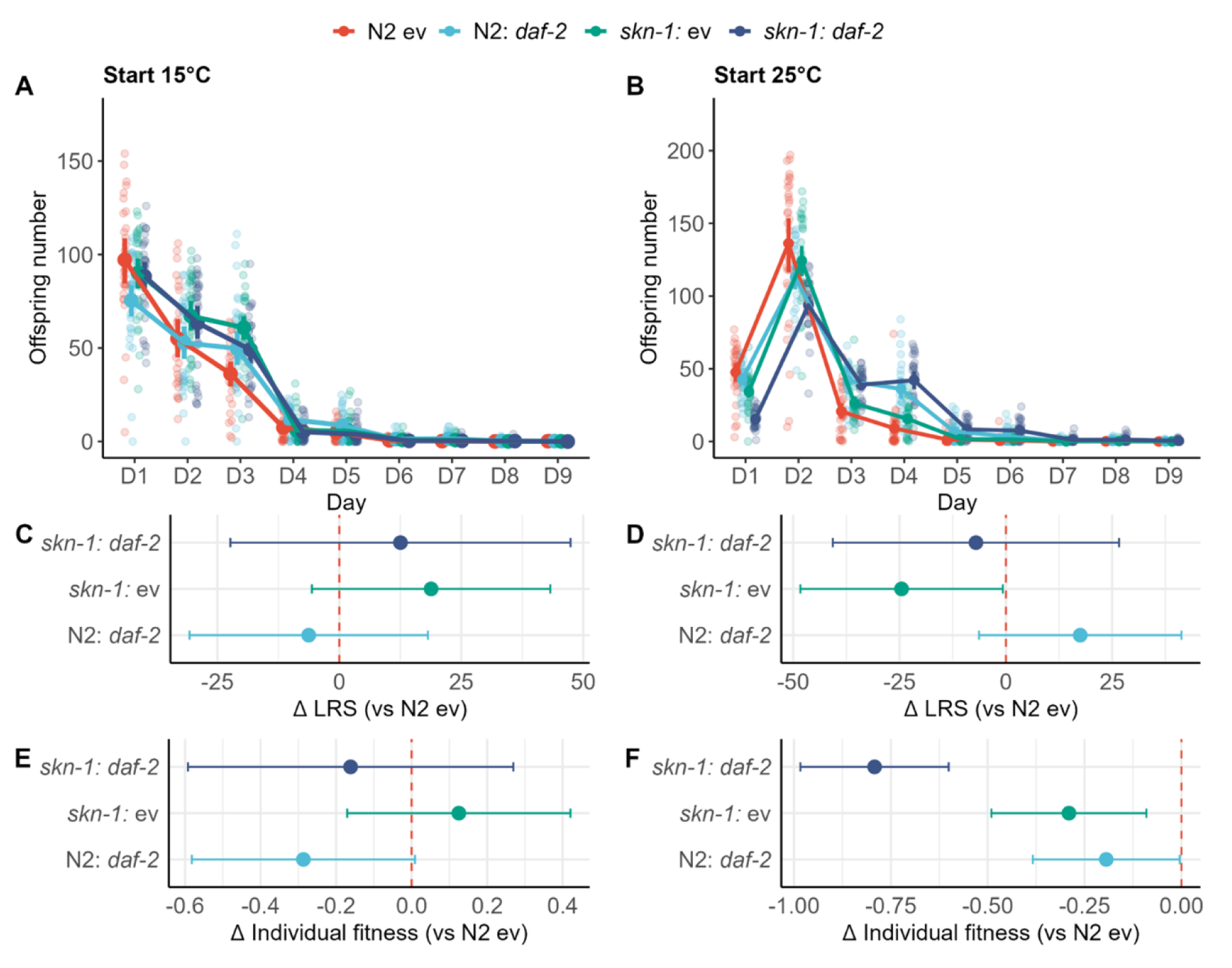
Developmental *daf-2* knockdown in *skn-1* mutants delays reproduction when starting development at 25°C, but not at 15°C under a 24-hour thermocycling regime, while N2s experience delayed reproduction following *daf-2* KD when starting thermocycling at 15°C. **(A, B)** Age-specific reproduction, **(C, D)** lifetime reproduction success (LRS), and **(E, F)** rate-sensitive individual fitness following *daf-2* knockdown or control treatment (empty vector plasmid) under a 24-hour rapid thermocycling regime between 15 and 25°C (n=30 per treatment x strain x temperature). Thermocycling temperatures began at 15°C **(A, C)** or 25°C **(B, D)** from the egg stage. In **(A, B)** points represent mean daily offspring production with 95% confidence intervals. In **(C - F)** coloured points represent mean differences (as compared to N2: ev controls) with 95% confidence intervals as calculated from linear models.

### 3.4. Temperature cycling at the L2 stage does not change ageing trajectories

While we found no effect of the developmental only thermocycle for *skn-1* mutants with *daf-2* knockdown, we did find that *daf-2* knockdown during development extended lifespan only in 15°C in both *skn-1* mutants and N2s (COXME: *skn-1* - Treatment: *Χ_1_^2^* = 25.555, *p* < 0.0001; N2 – Treatment: *Χ_1_^2^* = 12.230, *p* = 0.0005, Fig. 5A - C, Table S6). In 25°C, we saw no lifespan extension for any treatments in either strain (COXME: *skn-1* - Treatment: *Χ_1_^2^* = 0.063, *p* = 0.968; N2 – Treatment: *Χ_1_^2^* = 0.148, *p* = 0.700, Fig. 5D - F, Table S7). These effects were the same when matricides were included as natural deaths across both starting temperatures (Tables S6 & S7).

**Figure 5.**
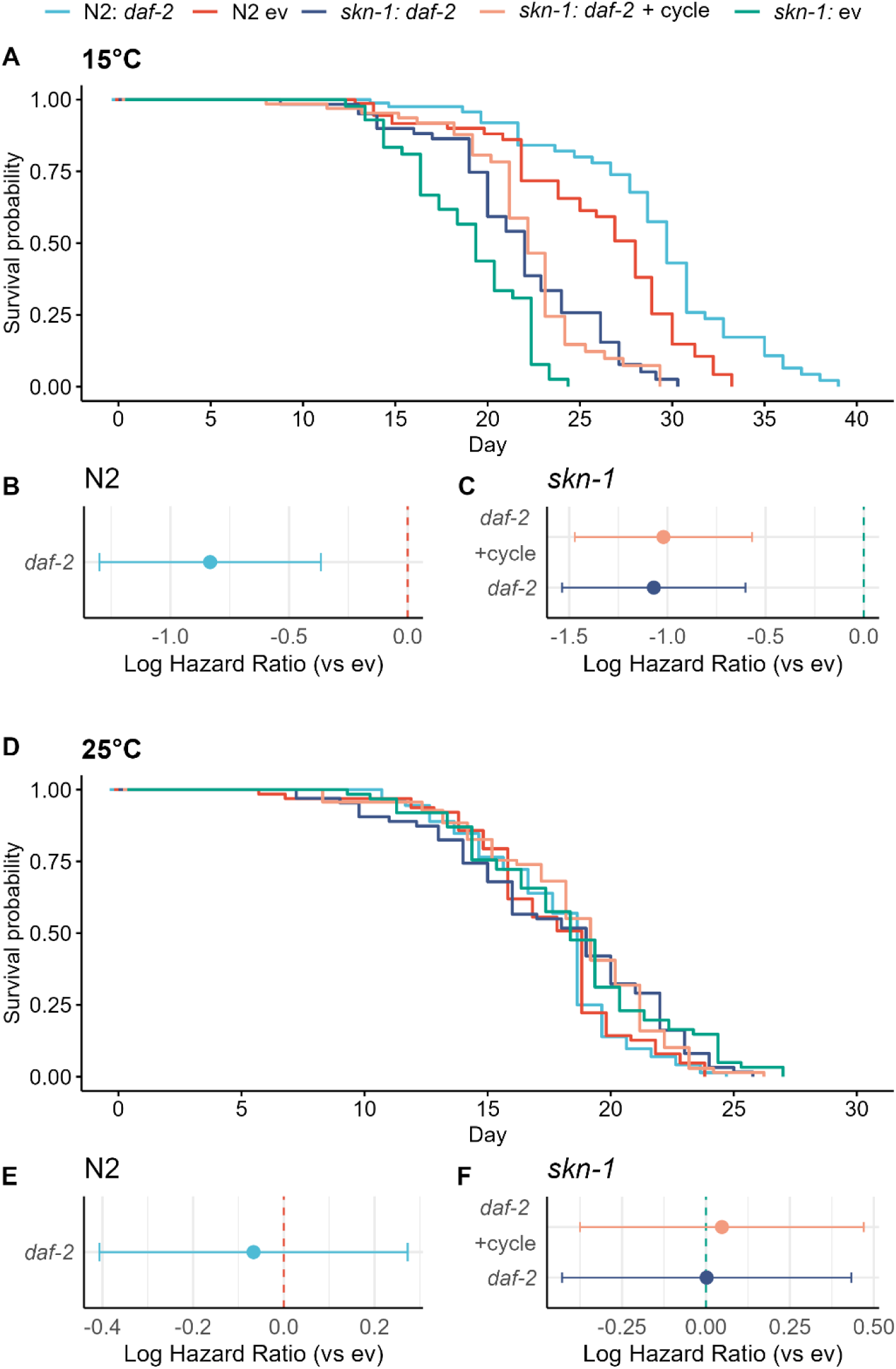
Developmental *daf-2* knockdown extends lifespan in N2 and *skn-1* mutants at 15°C even if *skn-1* individuals experience 25°C during L2 stage, but not at all at 25°C. **(A, D)** survival probability shown with Kaplan-Meier survival curves and **(B, C, E, F)** corresponding log hazard ratios under **(A - C)** 15°C (n > 44 per treatment x strain) and **(D - F)** 25°C (n > 62 per treatment x strain) following *daf-2* knockdown in N2 Bristol strain nematodes and *skn-1* loss of function mutants (with matricides excluded). Dotted lines represent baseline log hazard ratio set to zero for controls (N2 or *skn-1* with ev treatment). Log hazard ratios represented by means and 95% confidence intervals.

## 4. Discussion

It has long been thought that downregulating IIS through *daf-2* RNAi during development carries reproductive costs in *C. elegans* without extending lifespan (Dillin et al., 2002). Contrary to this, our work shows that developmental *daf-2* knockdown can extend lifespan at 15°C, but not at 25°C, suggesting strong age- and temperature-specific constraints to the late-life benefits of reduced IIS (rIIS). We also employed daily thermocycling between 15°C and 25°C to test the effects of developmental *daf-2* downregulation on lifespan and reproduction while shifting between temperatures with known opposing effects on ageing trajectories. Under these thermocycling regimes, developmental *daf-2* knockdown substantially extended lifespan. However, this effect depended on both *skn-1* function and the temperature at which thermocycling began, pointing to a crucial developmental window in which IIS, *skn-1*, and temperature interact to shape adult life-history trajectories.

While *daf-2* mediated lifespan extension persists in N2 and *skn-1* mutants at 15°C under constant conditions, it is lost in both strains at 25°C. Since 25°C approaches the upper thermal tolerance of *C. elegans* (with sterility occurring at 27°C), early-life exposure to this temperature likely represents a chronic, high-stress environment. Chronic exposure to high temperature is known to accelerate ageing and can overwhelm protective stress responses (Lithgow et al., 1995; Van Voorhies and Ward, 1999), potentially limiting the capacity of developmentally rIIS to extend lifespan under these conditions. In contrast, thermocycling may create an environment in which developmentally rIIS becomes beneficial by promoting stress-response priming rather than chronic adaptation to stress. Alternating exposure to 15°C and 25°C may allow transient activation of protective stress response pathways during high temperature exposure, followed by phases of recovery when temperatures are lower. This could potentially result in hormesis like effects. Under thermocycling, developmental *daf-2* knockdown may thus amplify the benefits of transient heat stress. This is consistent with evidence that mild or intermittent heat stress can extend lifespan, while chronic thermal stress shortens it (Olsen et al., 2006; Rea et al., 2005; Wu et al., 2009).

The timing of thermal stress exposure during development appears to be critical without *skn-1* functioning. Previous work has shown that transitions through the L1 and L2 larval stages are important for temperature-dependent life-history regulation (Zhang et al., 2015). This may be because the ‘decision’ to enter dauer is made primarily during the L1 and early L2 stages (Fielenbach and Antebi, 2008; Golden and Riddle, 1984), suggesting that life-history trajectories may be determined very early in life. However, shifting temperature specifically around the L2 stage had no effect on lifespan in otherwise constant conditions, suggesting that earlier or later stages may be more critical. Given the known role of *skn-1* during embryogenesis (Bowerman et al., 1992), we predict that differential temperature exposure during the egg stage may be particularly important for priming IIS-dependent stress responses that persist into adulthood.

Lifespan extension under thermocycling was associated with reduced and/or delayed reproductive output. Fitness declines under thermocycling may reflect a trade-off between increased stress responses to fluctuating conditions and reproductive investment. Notably, fitness costs were highest in *skn-1* mutants where thermocycling initiated at 25°C, possibly due to the engagement of alternative stress-resistance pathways. However, the strong age- and temperature-specificity of these effects suggests that they are not simply a direct resource allocation trade-off but may instead reflect a latent developmental program that mediates stress resistance and reproductive timing in response to early-life conditions. More broadly, the context-specificity we observe supports the idea that lifespan and ageing are condition-dependent traits (Chen and Maklakov, 2014; Williams et al., 2006), such that the environment in which an organism develops and lives can reshape senescence trajectories rather than these being fixed by a single allocation trade-off (Cohen et al., 2020).

The dual roles of *skn-1* in development and adult stress protection (An and Blackwell, 2003; Blackwell et al., 2015; Oliveira et al., 2009) imply that selection on *skn-1* activity is likely both highly age- and environment-dependent. Our results highlight how IIS plasticity during early development is integral to *C. elegans* life history, especially given its central role in regulating dauer entry and life-history strategies. Together, these findings support the existence of a critical developmental window during which IIS, *skn-1,* age, and environmental variability interact to shape ageing trajectories. Future work aimed at refining these methods and precisely defining the critical developmental windows indicated by our research could point to more specific constraints on developmental optimisation of IIS regulation for late-life survival.

## Acknowledgments

This work was supported by the European Research Council consolidator grant SELECTHAPLOID 101001341 awarded to SI and the Natural Environment Research Council grant NE/W001020/1 awarded to AAM.

## Author contributions

Conceptualization: SDI, SI, AAM

Formal analysis: SDI

Funding acquisition: SI, AAM

Investigation: SDI, AK

Methodology: SDI, AK

Supervision: SI, AAM

Visualization: SDI

Writing – original draft: SDI

Writing – review & editing: SDI, AK, SI, AAM

## Data availability

All code and data used for this study are available via figshare at https://doi.org/10.6084/m9.figshare.34529865.

