## Supplementary Tables for "Temperature and *skn-1* determine whether developmental *daf-2* knockdown extends *C. elegans* lifespan"

This file contains:

Tables S1 to S7

**Tables:**

**Table S1. Sample sizes and median lifespan for all *C. elegans* lifespan experiments at constant and cycling temperatures in N2 and *skn-1* mutants.** Sample sizes include total nematodes setup for assays (n_total_) and the number of individuals included in the final analysis, excluding matricides (n_censored_). Number of matricides (mat.) and ruptures causing expelling of intestines (exp.) also included. Any other censored individuals were lost either due to human error or nematodes dying due to crawling up the side of the agar plate. Median lifespan is calculated from the fit of Kaplan-Meier survival curves, along with lower (LCL) and upper (UCL) confidence limits produced by the same fit.

| *skn-1* mutants at constant 20˚C | | | | | | |
| --- | --- | --- | --- | --- | --- | --- |
| *Treatment* | *n_total_* | *n_censored_* | *mat.* | *exp.* | *median lifespan* | *LCL, UCL* |
| *daf-2* | 106 | 76 | 12 | 21 | 18 | 17, 19 |
| ev | 106 | 69 | 12 | 17 | 19 | 17, 19 |
| Thermocycling starting at 15˚C | | | | | | |
| ***skn-1*** *daf-2* | 96 | 79 | 17 | 0 | 19 | 18, 19 |
| ***skn-1*** ev | 96 | 71 | 25 | 0 | 19 | 19, 20 |
| ***N2*** *daf-2* | 90 | 58 | 32 | 0 | 29 | 28, 30 |
| ***N2*** ev | 96 | 73 | 23 | 0 | 21 | 20, 21 |
| Thermocycling starting at 25˚C | | | | | | |
| ***skn-1*** *daf-2* | 100 | 74 | 17 | 5 | 26 | 26, 27 |
| ***skn-1*** ev | 100 | 72 | 17 | 1 | 18 | 18, 18 |
| ***N2*** *daf-2* | 100 | 89 | 10 | 1 | 27 | 27, 30 |
| ***N2*** ev | 100 | 93 | 7 | 0 | 21 | 21, 21 |
| Constant 15˚C | | | | | | |
| ***skn-1*** *daf-2* | 87 | 42 | 23 | 19 | 20 | 20, 24 |
| ***skn-1*** *daf-2+L2* | 85 | 43 | 20 | 21 | 21 | 21, 23 |
| ***skn-1*** ev | 81 | 40 | 36 | 3 | 19 | 17, 21 |
| ***N2*** *daf-2* | 91 | 49 | 2 | 39 | 30 | 29, 31 |
| ***N2*** ev | 87 | 50 | 9 | 26 | 28 | 25, 29 |
| Constant 25˚C |  |  |  |  |  |  |
| ***skn-1*** *daf-2* | 67 | 62 | 0 | 5 | 19 | 16, 20 |
| ***skn-1*** *daf-2+L2* | 71 | 69 | 0 | 0 | 19 | 18, 20 |
| ***skn-1*** ev | 64 | 61 | 0 | 1 | 18 | 17, 19 |
| ***N2*** *daf-2* | 75 | 72 | 0 | 3 | 19 | 18, 19 |
| ***N2*** ev | 67 | 63 | 0 | 2 | 19 | 17, 19 |

**Table S2:** **Developmental *daf-2* knockdown does not affect age-specific reproduction in SKN-1 mutants at 20°C.** Model summary from mixed effects negative binomial generalised linear model with a zero-inflation parameter. ID is included as a random effect to control for repeated measures. *Χ^2^* from type III Anova is included to represent overall contribution of main effects and interactions.

|  | *Χ_1_^2^* | *Estimate* | *SE* | *z* | *p* |
| --- | --- | --- | --- | --- | --- |
| (Intercept) | 266.092 | 3.386 | 0.208 | 16.312 | <0.0001 |
| Treatment.ev | 0.720 | -0.107 | 0.126 | -0.849 | 0.396 |
| Day^2^ | 162.721 | -0.336 | 0.026 | -12.756 | <0.0001 |
| Day | 77.284 | 1.347 | 0.153 | 8.791 | <0.0001 |
| Treatment.ev*Day^2^ | 0.389 | -0.007 | 0.011 | -0.624 | 0.533 |
| Zero-inflation formula | | | | | |
| (Intercept) | | -6.778 | 1.120 | -6.050 | <0.0001 |
| Day | | 1.206 | 0.256 | 4.718 | <0.0001 |
| Random effects | | | | | |
| ID | | Variance: | <0.0001 | SD: | <0.0001 |

**Table S3. Developmental *daf-2* knockdown extends lifespan in N2s, but not SKN-1 mutants under a 24-hour thermocycling treatment that begins at 15°C.** Type II Anova summary from mixed effects cox proportional hazards model including contrasts (from emmeans() package) for treatments within strains.

| **15°C first** | | | | | | | | | | | | | | | |
| --- | --- | --- | --- | --- | --- | --- | --- | --- | --- | --- | --- | --- | --- | --- | --- |
| *Matricides included* | | | | | | | | | | | | | | | |
|  | *Χ_1_^2^* | *p* | Contrasts | | | | | | | | | | | | |
| Strain | 70.954 | <0.0001 |  | *Est.* | | | *SE* | | | *z* | | | *p* | | |
| Treatment | 45.512 | <0.0001 | N2 | | | | | | | | | | | | |
| Strain*Treatmt | 29.317 | <0.0001 | daf-ev | | -1.22 | | | | 0.18 | | | -6.75 | | | <0.0001 |
| Random effect: ID  Var: 0.009; SD < 0.0001 | | | SKN-1 | | | | | | | | | | | | |
|  |  |  | daf-ev | | | 0.06 | | 0.15 | | | 0.43 | | | 0.666 | |
| *Matricides censored* | | | | | | | | | | | | | | | |
|  | *Χ_1_^2^* | *p* | Contrasts | | | | | | | | | | | | |
| Strain | 134.444 | <0.0001 |  | *Est.* | | | *SE* | | | *z* | | | *p* | | |
| Treatment | 76.595 | <0.0001 | N2 | | | | | | | | | | | | |
| Strain*Treatmt | 61.563 | <0.0001 | daf-ev | | -2.10 | | | | 0.24 | | | -8.75 | | | <0.0001 |
| Random effect: ID  Var: 0.095; SD = 0.009 | | | SKN-1 | | | | | | | | | | | | |
|  |  |  | daf-ev | | | 0.30 | | 0.17 | | | 1.73 | | | 0.084 | |
| **25°C first** | | | | | | | | | | | | | | | |
| *Matricides included* | | | | | | | | | | | | | | | |
|  | *Χ_1_^2^* | *p* | Contrasts | | | | | | | | | | | | |
| Strain | 21.160 | <0.0001 |  | *Est.* | | | *SE* | | | *z* | | | *p* | | |
| Treatment | 94.447 | <0.0001 | N2 | | | | | | | | | | | | |
| Strain*Treatmt | 2.420 | 0.120 | daf-ev | | -1.69 | | | | 0.17 | | | -9.72 | | | <0.0001 |
| Random effect: ID  Var: <0.0001; SD = 0.004 | | | SKN-1 | | | | | | | | | | | | |
|  |  |  | daf-ev | | | -2.04 | | 0.19 | | | -10.91 | | | <0.0001 | |
| *Matricides censored* | | | | | | | | | | | | | | | |
|  | *Χ_1_^2^* | *p* | Contrasts | | | | | | | | | | | | |
| Strain | 17.702 | <0.0001 |  | *Est.* | | | *SE* | | | *z* | | | *p* | | |
| Treatment | 121.096 | <0.0001 | N2 | | | | | | | | | | | | |
| Strain*Treatmt | 7.324 | 0.007 | daf-ev | | -2.07 | | | | 0.19 | | | -11.00 | | | <0.0001 |
| Random effect: ID  Var: 0.004; SD <0.0001 | | | SKN-1 | | | | | | | | | | | | |
|  |  |  | daf-ev | | | -2.73 | | 0.22 | | | -12.64 | | | <0.0001 | |

**Table S4:** **Developmental *daf-2* knockdown does not affect age-specific reproduction in SKN-1 mutants under 24-hour thermocycling starting at 15°C but slows early reproduction in N2s.** **When starting at 25°C, *daf-2* knockdown slows down early reproduction in SKN-1 mutants and to a lesser extent, N2s.** Model summary from mixed effects negative binomial generalised linear model with a zero-inflation parameter, with estimates, standard errors, and z-values calculated for *daf-2* and SKN-1 compared to controls and N2. ID is included as a random effect to control for repeated measures. *Χ^2^* from type III Anova is included to represent overall contribution of main effects and interactions.

| **15°C first** | | | | | | | |
| --- | --- | --- | --- | --- | --- | --- | --- |
|  | | | Χ_1_^2^ | Estimate | SE | z | p |
| (Intercept) | | | 3914.828 | 5.269 | 0.084 | 62.57 | <0.0001 |
| Strain | | | 0.335 | 0.064 | 0.111 | 0.58 | 0.5627 |
| Treatment | | | 5.917 | -0.273 | -0.273 | -2.43 | 0.0150 |
| Day | | | 367.725 | -0.660 | -0.660 | -19.18 | <0.0001 |
| Strain*Treatment | | | 3.958 | 0.308 | 0.308 | 1.99 | 0.0467 |
| Strain*Day | | | 0.390 | 0.028 | 0.045 | 0.62 | 0.5324 |
| Treatment*Day | | | 14.570 | 0.160 | 0.042 | 3.82 | 0.0001 |
| Strain*Treatment*Day | | | 13.504 | -0.224 | 0.061 | -3.67 | 0.0002 |
| Zero-inflation formula | | | | | | | |
| (Intercept) | | | | -5.348 | 0.470 | -11.377 | <0.0001 |
| Day | | | | 0.708 | 0.083 | 8.494 | <0.0001 |
| Random effects | | | | | | | |
| ID | | | | Variance: | <0.0001 | SD: | <0.0001 |
| **25°C first** | | | | | | | |
|  | Χ_1_^2^ | | | Estimate | SE | z | p |
| (Intercept) | | 1468.867 | | 5.631 | 0.150 | 38.33 | <0.0001 |
| Strain | | 7.664 | | -0.504 | 0.182 | -2.77 | 0.006 |
| Treatment | | 7.111 | | 0.460 | 0.172 | 2.67 | 0.008 |
| Day | | 310.250 | | -0.645 | 0.037 | -17.61 | <0.0001 |
| Strain*Treatment | | 1.762 | | 0.188 | 0.142 | 1.33 | 0.184 |
| Strain*Day | | 11.249 | | 0.140 | 0.042 | 3.35 | 0.001 |
| Treatment*Day | | 49.505 | | -0.307 | 0.044 | -7.04 | <0.0001 |
| Zero-inflation formula | | | | | | | |
|  | | (Intercept) | | -47.309 | 22.711 | -2.083 | 0.037 |
| ns(Day, df = 3)1 | | | | 33.920 | 15.847 | 2.140 | 0.032 |
| ns(Day, df = 3)2 | | | | 86.232 | 42.453 | 2.031 | 0.042 |
| ns(Day, df = 3)3 | | | | 22.191 | 9.595 | 2.313 | 0.021 |
| Random effects | | | | | | | |
| ID | | Variance: | | <0.0001 | SD: | <0.0001 |  |

| **Table S5: Developmental *daf-2* knockdown does not affect lifetime reproductive success (LRS: total offspring produced), but SKN-1 mutants overall have lower LRS.** Type II Anova results for linear models of the additive effects of strain and treatment on LRS. Non-significant interactions terms between strain and treatment were removed to identify main effects. | | | | |
| --- | --- | --- | --- | --- |
| **15°C first** | | | | |
|  | *Sum squares* | *df* | *F* | *p* |
| Strain | 10441 | 1 | 3.392 | 0.068 |
| Treatment | 1163 | 1 | 0.378 | 0.540 |
| Residuals | 354035 | 115 |  |  |
| Residual SE: 55.48 on 115 df; Adjusted R^2^ 0.015 | | | | |
| F_2,115_ = 1.905, p = 0.154 | | | | |
| **25°C first** | | | | |
|  | *Sum squares* | *df* | *F* | *p* |
| Treatment | 9170 | 1 | 3.086 | 0.082 |
| Strain | 18032 | 1 | 6.069 | 0.015 |
| Residuals | 347646 | 117 |  |  |
| Residual SE: 54.51 on 117 df; Adjusted R^2^ 0.057 | | | | |
| F_2,117_ = 4.577, p = 0.012 | | | | |

**Table S6. Developmental *daf-2* knockdown extends lifespan in SKN-1 mutants and N2s at 15°C, with developmental temperature changes having no additional effect after *daf-2* knockdown in SKN-1 mutants.** Model summary from mixed effects cox proportional hazards model showing estimates for Hazard ratios (HR) and log Hazard ratios (LHR) and including *Χ^2^* from a type II Anova. Data including and excluding matricides were analysed separately. Analyses were performed within strain due to additional treatment for SKN-1.

| **SKN-1** | | | | | |
| --- | --- | --- | --- | --- | --- |
| **Matricides included** | | | | |  |
|  | *LHR coef* | *HR coef* | *SE (LHR)* | *z* | *p* |
| Tr_temp_daf | -0.912 | 0.402 | 0.220 | -4.14 | <0.0001 |
| Tr_temp_daf+cycle | -0.928 | 0.396 | 0.219 | -4.23 | <0.0001 |
| Random effect: | Plate.ID | Variance: | 0.077 | SD: | 0.277 |
| Anova (type II): Treatment_temp | | Χ_2_^2^ = 23.377 | | p = <0.0001 | |
| **Matricides censored** | | | | | |
|  | *LHR coef* | *HR coef* | *SE (LHR)* | *z* | *p* |
| Tr_temp_daf | -1.070 | 0.343 | 0.238 | -4.49 | <0.0001 |
| Tr_temp_daf+cycle | -1.020 | 0.361 | 0.230 | -4.43 | <0.0001 |
| Random effect: | Plate.ID | Variance: | 0.0004 | SD: | 0.020 |
| Anova (type II): Treatment_temp | | Χ_2_^2^ = 25.555 | | p = <0.0001 | |
| **N2** | | | | | |
| **Matricides included** | | | | | |
|  | *LHR coef* | *HR coef* | *SE (LHR)* | *z* | *p* |
| Treatment_daf-2 | -0.905 | 0.405 | 0.239 | -3.79 | 0.0001 |
| Random effect: | Plate.ID | Variance: | 0.064 | SD: | 0.252 |
| Anova (type II): Treatment_temp | | Χ_2_^2^ = 14.383 | | p = 0.0001 | |
| **Matricides censored** | | | | | |
|  | *LHR coef* | *HR coef* | *SE (LHR)* | *z* | *p* |
| Treatment_daf-2 | -0.833 | 0.435 | 0.238 | -3.50 | 0.0005 |
| Random effect: | Plate.ID | Variance: | 0.041 | SD: | 0.203 |
| Anova (type II): Treatment_temp | | Χ_2_^2^ = 12.23 | | p = 0.0005 | |

**Table S7. Developmental *daf-2* knockdown does not extend lifespan in SKN-1 mutants and N2s at 25°C, with developmental temperature changes having no effect after *daf-2* knockdown in SKN-1 mutants.** Model summary from mixed effects cox proportional hazards model showing estimates for Hazard ratios (HR) and log Hazard ratios (LHR) and including *Χ^2^* from a type II Anova. Data including and excluding matricides were analysed separately. Analyses were performed within strain due to additional treatment for SKN-1.

| **SKN-1** | | | | | |
| --- | --- | --- | --- | --- | --- |
| **Matricides included** |  |  |  |  |  |
|  | *LHR coef* | *HR coef* | *SE (LHR)* | *z* | *p* |
| Tr_temp_daf_25 | -0.004 | 0.996 | 0.180 | -0.02 | 0.981 |
| Tr_temp_daf_25+cycle | -0.148 | 0.862 | 0.184 | -0.81 | 0.420 |
| Random effect: | Plate.ID | Variance: | 0.048 | SD: | 0.219 |
| Anova (type II): Treatment_temp | | Χ_2_^2^ = 0.845 | | p = 0.656 | |
| **Matricides censored** |  |  |  |  |  |
|  | *LHR coef* | *HR coef* | *SE (LHR)* | *z* | *p* |
| Tr_temp_daf_25 | 0.002 | 1.002 | 0.220 | 0.01 | 0.993 |
| Tr_temp_daf_25+cycle | 0.048 | 1.049 | 0.216 | 0.22 | 0.825 |
| Random effect: | Plate.ID | Variance: | 0.063 | SD: | 0.251 |
| Anova (type II): Treatment_temp | | Χ_2_^2^ = 0.065 | | p = 0.968 | |
| **N2** | | | | | |
| **Matricides included** |  |  |  |  |  |
|  | *LHR coef* | *HR coef* | *SE (LHR)* | *z* | *p* |
| Treatment_daf-2 | -0.173 | 0.841 | 0.179 | -0.97 | 0.334 |
| Random effect: | Plate.ID | Variance: | 0.049 | SD: | 0.222 |
| Anova (type II): Treatment_temp | | Χ_2_^2^ = 0.934 | | p = 0.334 | |
| **Matricides censored** |  |  |  |  |  |
|  | *LHR coef* | *HR coef* | *SE (LHR)* | *z* | *p* |
| Treatment_daf-2 | -0.067 | 0.9335 | 0.173 | -0.38 | 0.700 |
| Random effect: | Plate.ID | Variance: | <0.0001 | SD: | 0.009 |
| Anova (type II): Treatment_temp | | Χ_2_^2^ = 0.148 | | p = 0.700 | |
